# NLR immune receptors can exhibit tissue-specific expression patterns across legume species

**DOI:** 10.64898/2026.01.25.701577

**Authors:** Rita Maravilha Marques, Carmen Santos, Hsuan Pai, Maria Carlota Vaz Patto, Sophien Kamoun, Jiorgos Kourelis

## Abstract

Pathogen pressure threatens legume crop productivity worldwide. Nucleotide-binding leucine-rich repeat (NLR) immune receptors serve as crucial plant resistance genes, recognizing pathogens and triggering immunity. However, the extent and patterns of NLR expression in different tissues and organs, notably across evolutionary time, remain largely uncharacterized. To investigate tissue-specificity of NLR expression in the Fabaceae (legumes), we conducted comparative analyses integrating phylogenomics and transcriptomics in root and shoot tissues across different legume species. The NLR repertoires of 28 legumes were grouped into five monophyletic clades: coiled-coil NLR (CC-NLR), Toll/interleukin-1 receptor NLR (TIR-NLR), G10-subclade CC NLR (CC_G10_-NLR), RESISTANCE TO POWDERY MILDEW 8-like CC NLR (CC_R_-NLR), and TIR-NB-ARC-like β-propeller WD40/tetratricopeptide repeats (TNPs). Most legume NLRs belonged to CC-NLR and TIR-NLR clades, followed by CC_G10_-NLR, CC_R_-NLR, and TNP clades. In seven of these species, comparative analysis of NLR expression in leaves versus roots revealed that over half (∼57%) of expressed NLR genes showed predominant expression in one tissue: 34% in roots (451/1336), and 23% in leaves (311/1336). We identified 324 root-specific NLRs, 171 leaf-specific NLRs, and 841 non-specific NLRs, with an average tissue specificity per species of 32%. The closely related species grass pea (*Lathyrus sativus*) and pea (*Pisum sativum*) were an exception, showing higher levels of leaf-specific rather than root-specific NLR expression. We also identified conserved tissue expression patterns across legume species, resulting in a comprehensive resource describing tissue expression bias, enrichment, and specificity for 113 phylogenetic NLR subclasses. These legume NLR repertoires will support comparative studies between species and inform precision-breeding programs considering tissue expression patterns.

## INTRODUCTION

The legume family comprises 765 genera and over 19,600 species, representing one of the largest and most diverse plant families (Cannon *et al*., 2015; Wojciechowski, 2019). Annual production reaches 445 million tons of legume feed and fodder globally (*FAOSTAT*, 2022), with human consumption of pulses amounting to 101 million tons in 2024 (OECD/FAO, 2025). Legumes contain high protein levels (16-50%) and fix atmospheric nitrogen through symbiotic relationships, making them agriculturally and economically important while enriching soil fertility (Cannon *et al*., 2015). However, diseases including mildews, rusts, blight, molds, rots, and wilts constrain legume productivity and threaten global food security, with agricultural losses from diseases reaching 45% globally (Rubiales *et al*., 2015; Dutta *et al*., 2022; Zhou *et al*., 2025). Chemical control remains a widely used method for disease management. However, its economic, environmental, and public health disadvantages highlight the urgent need for more sustainable pest management strategies (Zhou *et al*., 2025). Resistant plant varieties represent the most efficient, cost-effective, and environmentally sustainable approach to disease management (Rubiales *et al*., 2015). Understanding plant immunity and pathogen defense mechanisms is crucial for improving resistance breeding efficacy (Bentham *et al*., 2020).

Nucleotide-binding leucine-rich repeat proteins (NLRs) are among the most extensively studied and economically relevant gene families due to their importance in disease resistance (van Wersch *et al*., 2020). NLRs act as immune receptors, functioning in immunity by directly and indirectly detecting pathogen proteins (Förderer and Kourelis, 2023). In the absence of recognition, these pathogen proteins, known as effectors, manipulate the host, allowing the pathogen to colonize and cause disease (Khan *et al*., 2018).

Canonical plant NLRs consist of a variable N-terminal domain, a central NB-ARC module (for Nucleotide-Binding APAF-1, certain plant *R* gene products, and CED-4), and a C-terminal Leucine Rich Repeat (LRR) domain (Van Ooijen *et al*., 2008). The NB-ARC module consists of three domains: a nucleotide-binding domain (NBD), a Helical Domain 1 (HD1), and a Winged-Helix Domain (WHD). This module contains a functional ATPase and regulates NLR activity through its nucleotide-binding state (Van Ooijen *et al*., 2008). As the most conserved part of an NLR, this NB-ARC module is useful for determining evolutionary relationships between NLRs through global alignments (Kourelis *et al*., 2021). The LRR domain is highly variable and functions in both direct and indirect pathogen effector detection, stabilizing the active and inactive states of the NLR (Förderer *et al*., 2022). Beyond the canonical domains, some plant NLRs have additional “integrated domains” which play major roles in effector recognition (Cesari *et al*., 2014; Białas *et al*., 2021).

Direct and indirect pathogen effector detection by the LRR results in a conformational switch in the NB-ARC module, allowing exchange of ADP and ATP. This results in homo-oligomerization of the NLR into either tetrameric, pentameric, or hexameric complexes known as “resistosomes” (Förderer *et al*., 2022). Resistosome formation activates the N-terminal domain, initiating immune signaling. Alternatively, some NLRs form pairs and networks, where one NLR is required for pathogen effector detection (sensor-NLR), while another is required for immune signaling (helper-NLR) (Adachi, Derevnina and Kamoun, 2019). Overall, the emerging view is to move away from a one-size-fits-all model of NLR structure and function and to appreciate the remarkable diversity of these proteins (Kourelis *et al*., 2021; Contreras *et al*., 2023).

Based on phylogenetic trees of the NB-ARC module, flowering plant NLRs have been divided into four major monophyletic classes. These classes are named after their most common N-terminal domains, though individual members may lack these domains or carry non-canonical integrated domains (Barragan *et al*., 2021; Białas *et al*., 2021). N-terminal domains include three sequence-divergent but structurally conserved 4-helical bundle (4HB) domains: the Rx-type coiled-coil (CC) domain, RESISTANCE TO POWDERY MILDEW 8 (RPW8)-type CC (CC_R_), or the “autonomous NLR clade”-type CC (CC_G10_), or a Toll/Interleukin-1 Receptor/Resistance (TIR) domain (Förderer and Kourelis, 2023). Upon activation, these canonical N-terminal domains assemble into higher-order complexes that mediate downstream immune signaling (Wang *et al*., 2019; Förderer *et al*., 2022). CC-domain-containing NLRs (CC-NLRs) are known to form pentameric or hexameric resistosomes, where the first α helix of the CC-domain forms a funnel that acts as a calcium (Ca^2+^)-permeable nonselective cation pore at the plasma membrane (Wang *et al*., 2019; Bi *et al*., 2021). Similarly, CC_G10_- and CC_R_-NLRs oligomerize upon activation and act as Ca^2+^ permeable pores (Jacob *et al*., 2021; Lee *et al*., 2021; Förderer and Kourelis, 2023). Activated CC_R_-NLRs localize not only to the plasma membrane but also to organellar membranes, including the endoplasmic reticulum, chloroplast, and mitochondria (Ibrahim *et al*., 2023). TIR-domain-containing NLRs (TIR-NLRs) form tetrameric resistosomes upon activation, where the TIR domain displays NAD+ hydrolase activity, producing signals that lead to heterodimerization of the lipase-like protein ENHANCED DISEASE SUSCEPTIBILITY 1 (EDS1) with either SENESCENCE ASSOCIATED GENE 101 (SAG101) or PHYTOALEXIN DEFICIENT 4 (PAD4) (Baggs *et al*., 2020; Pruitt *et al*., 2021). These, in turn, activate members of the CC_R_-NLRs N-REQUIREMENT GENE 1 (NRG1) or ACTIVATED DISEASE RESISTANCE 1 (ADR1) subfamilies, respectively (Dong *et al*., 2016; Castel *et al*., 2019; Sun *et al*., 2021).

In addition to the four NLR classes found in flowering plants, a fifth small, highly conserved monophyletic class called TIR-NB-ARC-like-β-propeller WD40/TPRs (TNPs) exists, which phylogenetically clusters with non-plant NLRs (Kourelis *et al*., 2021). These TNPs contain a phylogenetically distinct TIR domain acquired independently from TIR-NLRs (Johanndrees *et al*., 2023). TNPs lack the LRR of canonical plant NLRs and instead contain WD40 amino acid repeats and tetratricopeptide-like repeats (TPRs), which could serve as a ligand-binding domain (Johanndrees *et al*., 2023).

Because of their effector detection role, the NLR family tends to be among the most polymorphic in plant genomes, generating substantial variations in NLR sequence diversity and copy number (Zheng *et al*., 2016; Kourelis *et al*., 2021). Current NLR knowledge remains largely confined to a few model and crop species such as *Arabidopsis thaliana*, Solanaceae, and cereals, despite the agricultural significance of Fabaceae crops (Kourelis *et al*., 2021; Contreras *et al*., 2023; Förderer and Kourelis, 2023). The complete set of NLRs within a genome (the NLRome) of certain legume species has been recently described, including soybean (*Glycine max*), common bean (*Phaseolus vulgaris*), barrel medic (*Medicago truncatula*), peanut (*Arachis hypogaea*), red clover (*Trifolium pratense*), *Lotus japonicus*, pea (*Pisum sativum*), grass pea (*Lathyrus sativus*), lentil (*Lens culinaris*), faba bean (*Vicia faba*), common vetch (*Vicia sativa*), adzuki bean (*Vigna angularis*), mung bean (*Vigna radiata*), cowpea (*Vigna unguiculata*), and narrowleaf lupin (*Lupinus angustifolius*), among other species (Qureshi *et al*., 2023; Rani *et al*., 2023; Negi *et al*., 2024; Sultan *et al*., 2024). Variation in legume NLR abundance is remarkable, even within the same genus, with NLRome size varying between 77 and 348 NLRs among *Glycine* species (Sultan *et al*., 2024).

Variation in NLR expression across tissues and organs is a surprisingly understudied aspect of the plant immune system (Lüdke *et al*., 2023). Precise regulation is critical, as the misexpression of NLRs can lead to autoimmunity and significant fitness costs, underscoring the evolutionary pressure to confine their expression to where it is most needed (Palma *et al*., 2010; Richard *et al*., 2018; Fick *et al*., 2022). Roots and leaves are physiologically distinct and have fundamentally different microenvironments (Bodenhausen *et al*., 2013; Munch *et al*., 2017). Additionally, roots and leaves are exposed to different types and levels of pathogens, pests, and beneficial microbes. Interestingly, *Arabidopsis thaliana* shows predominant leaf NLR expression, while the legume *L. japonicus* shows an opposite trend towards root expression, highlighting potential differences between these species in pathogen exposure patterns (Munch *et al*., 2018). In other species, some NLRs display tissue-specificity. For example, in tomato, the helper-NLR NLR-REQUIRED FOR CELL-DEATH (NRC) 6 and the sensor-NLRs Hero and MeR1, which provide immunity towards cyst- and root-knot nematodes, respectively, are root-specifically expressed (Lüdke *et al*., 2025).

Here, we integrated phylogenomic classification with transcriptomic profiling to characterize NLR tissue expression patterns across diverse legume species, generating a community resource that supports future comparative studies. We first identified and characterized the NLRomes of 28 legume species using NLRtracker and classified them into five monophyletic classes using the RefPlantNLR dataset as reference (Kourelis *et al*., 2021). We then conducted comparative transcriptomic analysis of NLR expression in leaves and roots for seven of these species. Our study revealed that legumes show predominant NLR gene expression in roots, although both grass pea (*Lathyrus sativus*) and pea (*P. sativum*) displayed higher rates of leaf-specific expression. We investigated whether tissue specificity varied according to monophyletic NLR categories, revealing enhanced tissue-specificity compared to other gene families. Additionally, we observed tissue expression patterns that were conserved across species. Our findings offer insight into how spatial expression contributes to NLR function and evolution, highlighting the role of selective pressures in shaping the deployment of immune receptors in plant tissues.

## MATERIAL AND METHODS

### Species selection and genome acquisition

We analyzed the predicted proteomes of 28 legume species representing six major clades: Hologalegina, Phaseoloid, Dalbergioids, Genistoids, Mimosoids, and Caesalpinioideae (**Table S1**). Four outgroup species with well-characterized NLRomes were included for comparative analysis: *Arabidopsis thaliana* (Ecotype Col-0; GCF_000001735.4; Annotation by TAIR and Araport), *Solanum lycopersicum* (cv. Heinz 1706; GCF_000188115.5; NCBI Solanum lycopersicum Annotation Release 103), *Oryza sativa* (Japonica Group, strain Nipponbare; GCF_001433935.1; NCBI Oryza sativa Japonica Group Annotation Release 102), and *Zea mays* (cv. B73; GCF_902167145.1; NCBI Zea mays Annotation Release GCF_902167145.1-RS_2025_02).

Genome assemblies and proteomes were obtained from multiple repositories. From the GigaScience Database (GigaDB): *Vicia sativa* (wild accession from South Australia, GCA_021764765.1) and *Cercis canadensis* (wild accession from Iowa, United States of America, GCA_003255065.1). From the Legume Information System (LIS): *Lotus japonicus* (ecotype Gifu, GCA_012489685.2), *Lupinus albus* (cultivar Amiga, GCA_009771035.1), *Lupinus angustifolius* (genotype Tanjil, GCA_001865875.1), *Phaseolus acutifolius* (wild accession W6 15578, *Phaseolus acutifolius* WLD v2.0), *Phaseolus lunatus* (accession G27455, bioproject PRJNA596114), *Vigna angularis* (genotype Gyeongwon IT213134, GCA_000465365.1), and *Vigna radiata* (genotype VC1973A, GCA_000741045.2). *Prosopis cineraria* (wild accession growing in 24°17′18.3″ N 55°43′36.2″ E) was obtained from Zenodo (https://zenodo.org/record/6720540). *Lens culinaris* genome assembly (CDC Redberry; Lcu.2RBY) was obtained from the authors of Haile *et al*. (2021). The remaining genomes were retrieved from the National Center for Biotechnology Information (NCBI) platform. Complete source information and genome quality metrics are provided in **Table S1**.

### Phylogenetic analysis of legume species

Phylogenetic relationships among analyzed species were determined using three chloroplast genes: ATP synthase subunit alpha (*atpA*), maturase K (*matK*), and the large subunit of ribulose bisphosphate carboxylase (*rbcL*) (Ohashi, Tateishi and Donovan Bailey, 2001; Chebet *et al*., 2022). Gene sequences were aligned using Clustal Omega v1.2.2 in Geneious Prime v2022.2.2 (https://www.geneious.com) with default parameters (group sequences by similarity, use fast clustering with mBed algorithm, and mBed guide trees cluster size of 100). A consensus phylogenetic tree for 28 legume species was constructed using the Geneious Tree Builder with the Tamura-Nei neighbor-joining method, *Z. mays* as outgroup, and 10,000 bootstrap resamples.

### Genome quality assessment

Genome quality was evaluated using two metrics: contiguity and completeness. Contiguity was assessed using contig N50 size, with genomes having N50 ≥ 1 Mbp considered high-quality (Astashyn *et al*., 2024). Completeness was evaluated using Benchmarking Universal Single-Copy Orthologs (BUSCO v5; Manni *et al*., 2021) with fabales_odb as reference for legumes and embryophyta_odb for outgroup species (**Table S2**). Only proteomes with <5% missing core orthologs were considered complete and retained for further analysis.

### NLR identification and classification

NLRs were extracted from each proteome using NLRtracker v1.3.1 (Kourelis *et al*., 2021). For accurate class assignment, NLRtracker NB-ARC output files were aligned with RefPlantNLR extracted NB-ARC sequences from supplemental datasets 9, 13, and TN_OTHER (Kourelis *et al*., 2021).

Sequence alignments were performed using Clustal Omega v1.2.2 (Sievers and Higgins, 2014) with default parameters in Geneious Prime v2025.2.1 (https://www.geneious.com). Approximately maximum likelihood phylogenetic trees were generated using FastTree v2.1.11 (Price *et al*., 2010) with default settings and pseudo-count settings for highly gapped sequences. Trees were rooted using non-plant NLRs from the RefPlantNLR dataset (Kourelis *et al*., 2021).

Phylogenetic trees were visualized using iTOL (Letunic and Bork, 2024). NLR classes were assigned based on monophyletic clustering with respective reference NLRs from RefPlantNLR (Kourelis *et al*., 2021). For example, CC_R_-NLRs were identified by clustering with NRG1 and ADR1 reference sequences. Complete classification details are provided in **Table S3**.

Alternative splicing variants were identified through genome localization analysis and removed to count only one NLR per locus (**Table S4**). NLR abundance was normalized by dividing the total NLR count by the number of protein-coding loci in each genome (Sekhwal *et al*., 2015) (**Tables S4**, **S5**). Correlation between proteome size and NLR number was assessed using Pearson’s correlation coefficient. Statistical analyses were performed in R using custom scripts available at https://github.com/MaravilhaRM/NLR_Classes_MaravilhaR.git and Maravilha Marques *et al*., 2026.

### Plant material and growth conditions

For tissue expression analysis, we selected seven closely related species from Hologalegina and Phaseoloid clades with their corresponding genome assemblies and annotations: *Lathyrus sativus* LS007 (JIC_Lsat_v2.1.1; GCA_963859935.3), *Pisum sativum* Caméor (CAAS_Psat_ZW6_1.0; GCA_024323335.2), *Lens culinaris* CDC Redberry (Lcu.2RBY), *Cicer arietinum* kabuli variety (genotype CDC Frontier; GCA_000331145.1), *Medicago truncatula* A17 (GCF_003473485.1_MtrunA17r5.0-ANR-EGN-r1.9; NCBI Medicago truncatula Annotation Release GCF_003473485.1-RS_2025_04), *Glycine max* Williams 82 (Wm82.a4.v1; GCA_000004515.5), and *Phaseolus vulgaris* genotype G19833 (phavu.G19833.gnm2.ann1.PB8d; GCA_000499845.1). Plants were cultivated in 0.5 L pots containing a mixture of 50% soil, 25% vermiculite, 25% peat in controlled environment chambers at 22°C with a 12/12-hour photoperiod.

### RNA extraction and sequencing

Leaf and root tissues were harvested separately from plants with three fully expanded leaves, immediately frozen in liquid nitrogen, and stored at -80°C until processing. Plants showed spontaneous nodulation upon harvest. Total RNA was extracted from approximately 100 mg of tissue using the GeneJET Plant RNA Purification Mini Kit (Thermo Scientific, Vilnius, Lithuania) according to manufacturer’s protocols, with three biological replicates per tissue and species.

RNA samples were treated with Turbo DNase I (Ambion, Austin, TX, USA) following manufacturer’s instructions. RNA concentration was quantified using a Qubit 2.0 Fluorometer (Invitrogen, Life Technologies, CA, USA) with Qubit RNA BR Assay Kit. RNA purity was assessed by measuring absorbance ratios at 260/280 nm and 260/230 nm using a NanoDrop 2000c spectrophotometer (Thermo Scientific, Passau, Germany). RNA integrity was verified by electrophoresis on 1% agarose gels stained with SYBR Safe (Life Technologies, CA, USA) at 100 V for 30 minutes.

RNA samples were submitted for Illumina NovaSeq 6000 Sequencing PE150 (40 million paired end reads per sample) at Novogene (UK) Co. Raw sequence data quality was assessed using FastQC (Andrews, 2010). Read preprocessing included removal of the first 10 base pairs corresponding to barcode contamination, removal of Illumina Universal adapters, and trimming of bases with Phred quality scores <20 using cutadapt v4.0 (Martin, 2011).

### RNA-seq data analysis

Processed reads were aligned to the corresponding reference genomes used for NLR discovery using HISAT2 v2.2.1 (Kim *et al*., 2015). SAMtools v1.9 (Li *et al*., 2009) was used for BAM file sorting and conversion. Transcript quantification was performed using StringTie v2.2.1 (Pertea *et al*., 2016). Genes with average expression <5 transcripts across all samples and alternative splicing variants were excluded from analysis.

Count data normalization was performed using the trimmed mean of M-values (TMM) method (**Table S6**). Tissue specificity assessment involved averaging expression values across leaf and root tissue replicates. Genes were classified as tissue-specific if they showed significant expression (average TMM >0.5 across three replicates) in one tissue while remaining lowly expressed (average TMM <0.5) in the other tissue (**Table S7**).

Differential expression analysis was conducted in R using DESeq2 with root samples as the reference (Anders and Huber, 2010; Love *et al*., 2014). Differentially expressed genes (DEGs) between leaf and root tissues were identified using a false discovery rate threshold of 0.05 and a log_2_ fold change |>1| (**Table S8**).

### Phylogenetic analysis of NLR subclasses

Phylogenetic analyses of expressed NLRs were performed using a combination of Geneious Prime v2025.2.1 with Clustal Omega alignment tool v1.2.2 (Sievers and Higgins, 2014) and FastTree plugin v2.1.11 (Price *et al*., 2010). NB-ARC domains from the expressed NLRs were aligned with the RefPlantNLR extracted NB-ARC domain dataset and phylogenetic trees constructed as described above to investigate clustering patterns with reference NLRs. NLR subclasses were extracted based on branch distances (0.315 for TNPs, 1.22 for CC_G10_-NLRs, 1.25 for CC_R_-NLRs, 1.2 for CC-NLRs, and 1.24 for TIR-NLRs) per class subtree (**Table S9**). NLR subclass characteristics (number of NLRs, number of species present, and different expression classes) were summarized in **Table S10**.

## RESULTS

### Legume phylogeny and genome quality assessment

Accurate phylogenetic relationships and high-quality genomic resources are essential for comparative NLR analysis. To establish evolutionary relationships among legume species and assess resource quality, we analyzed three chloroplast genes and evaluated genome assembly metrics. Phylogenetic analysis of three chloroplast genes (*atpA*, *matK*, and *rbcL*) confirmed expected evolutionary relationships among the 28 legume species and 4 outgroups (**Figure S1A**). Both monocots (*Z. mays* and *O. sativa*) clustered together, followed by the dicot outgroups (*S. lycopersicum* and *A. thaliana*), then the mimosoids (*C. canadensis* and *P. cineraria*), and finally the papilionoid clade species (**Figure S1A**).

Genome quality assessment revealed substantial variation in assembly contiguity and completeness across legume resources (**Figure S1B, C; Tables S1, S2**). Fifteen of 28 legume genomes were highly contiguous (contig N50 > 1 Mbp), while 16 predicted proteomes were considered complete (<5% missing BUSCOs) (**Figure S1C; Table S2**). Duplicated BUSCO scores ranged widely from 1.6% in *C. arietinum* to 95.2% in *A. hypogaea*, highlighting significant variation in genome duplication rates across legume species (**Figure S1C**). Only 11 of the 32 predicted proteomes met both contiguity and completeness criteria, though all predicted proteomes were retained for subsequent analyses.

These analyses provided the phylogenetic framework for comparative studies and revealed substantial variation in genomic resource quality across the legume family.

### Legume NLR gene counts exhibit extensive diversity and phylogenetic structure

Predicted NLR repertoires vary dramatically across plant species and families. To characterize legume NLR diversity and classify them into functional groups, we identified NLRs across 28 legume species and assigned them to monophyletic classes using phylogenetic analysis.

Legume NLR gene counts range from 88 in *L. albus* to 970 in *A. hypogaea* (**Figure 1A**). A moderate positive correlation (R = 0.59, p-value < 0.001) exists between NLR gene counts and predicted proteome size (**Figure 1A**). The proportion of predicted protein-coding loci encoding NLRs varied widely, from 0.12% in *L. albus* to 2.27% in *P. alba* (**Figure 1B**). Three legume species (*M. truncatula*, *A. hypogaea*, and *P. alba*) contained up to three times as many NLRs as expected for their predicted proteome size, harboring 645 to 931 predicted NLRs (**Figure 1B**). These species contrasted sharply with *C. arietinum*, *V. angularis*, *Aeschynomene evenia*, *P. sativum*, *P. lunatus*, *C. cajan*, *L. albus*, and *L. angustifolius*, which had fewer predicted NLRs than expected (**Figure 1B**).

**Figure 1.**
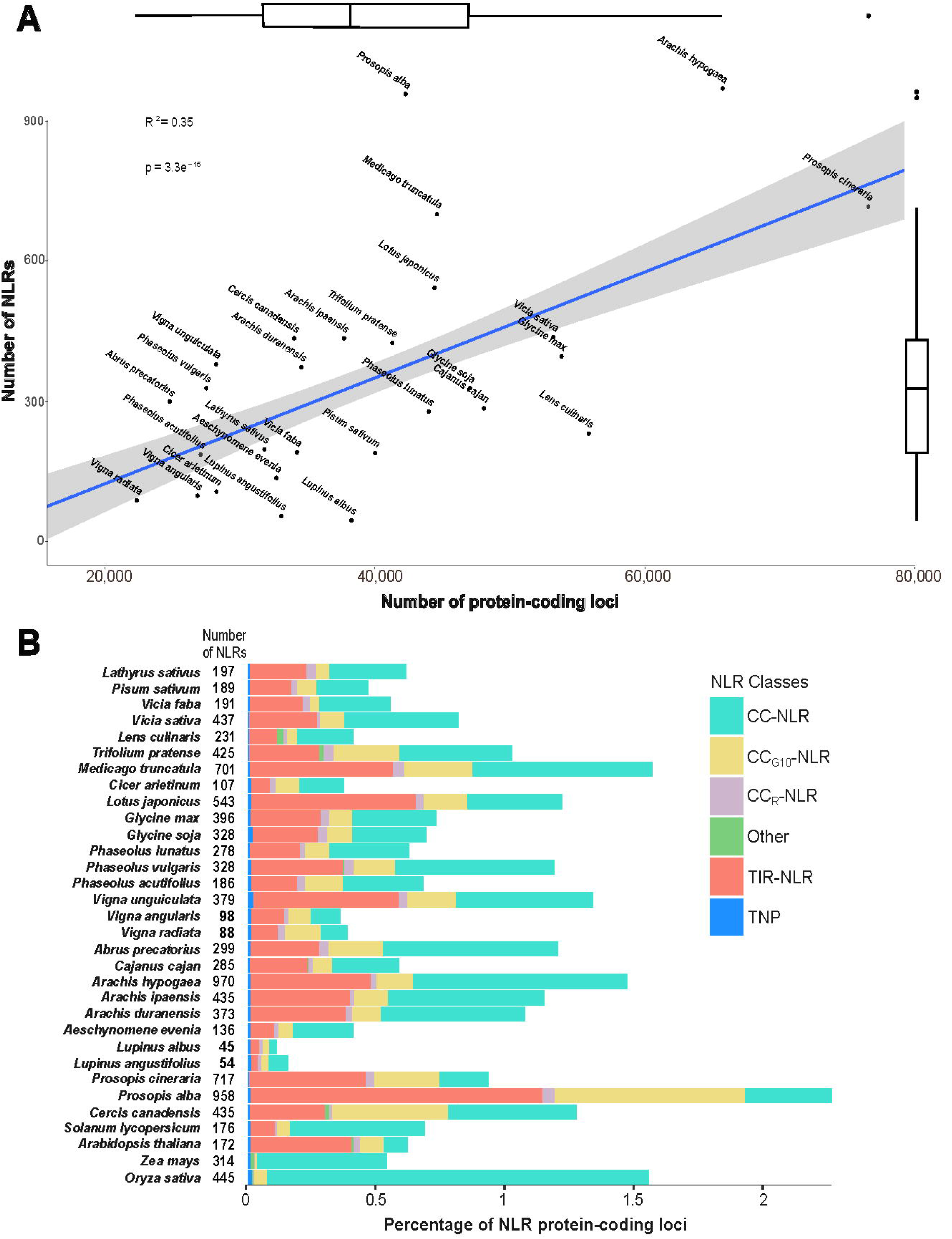
Legume NLR gene loci and counts. (**A**) NLR gene counts correlate moderately with predicted proteome size across legume species. Scatter plot showing NLR gene number versus protein-coding loci for each species. (**B**) NLR class distribution varies across legume phylogeny. Percentage of NLR-coding loci for each legume and outgroup species, arranged by phylogenetic relationships. Numbers indicate total NLR-coding protein loci per genome. Color coding: turquoise (CC-NLRs), yellow (CC_G10_-NLRs), lilac (CC_R_-NLRs), salmon (TIR-NLRs), blue (TNPs), and green (NLRs that did not cluster with monophyletic clades present in the RefPlantNLR dataset).

NLR classification through NB-ARC domain alignment with RefPlantNLR dataset revealed clustering into canonical flowering plant NLR monophyletic clades. CC-NLRs predominated across species, followed closely by TIR-NLRs, then CC_G10_-NLRs, with CC_R_-NLRs and TNPs being least abundant (**Figure 1B, S2**). Six species (*O. sativa*, *Z. mays*, *A. thaliana*, *P. vulgaris*, *T. pratense*, and *L. culinaris*) contained NLR groups that failed to cluster with reference NLRs and were classified as “Other” (**Figure 1B**). NLR class distribution largely reflected interspecific phylogenetic relationships, with legume NLR profiles showing greater similarity to dicot outgroups than to monocots (**Figure 1B**). Monocot outgroups (*O. sativa* and *Z. mays*) completely lacked TIR-NLRs. *V. radiata* and the early-diverging legumes *P. cineraria*, *P. alba*, and *C. canadensis* showed overrepresentation of CC_G10_-NLRs relative to other species (**Figure 1B**).

Legume NLRomes exhibit extensive diversity in gene counts and phylogenetic structure, with class distributions reflecting evolutionary relationships.

### Tissue identity accounts for greater expression variation than species differences

Understanding the relative contributions of tissue type versus species identity to gene expression patterns is crucial for interpreting tissue expression analyses. To determine the primary sources of expression variation in legumes, we performed transcriptomic analysis across seven species in two tissue types. It is noteworthy that these seedlings showed root nodules upon harvest, regardless of species.

Transcriptomic analysis of BUSCO fabales orthologues across leaf and root tissues revealed tissue as the primary source of expression variation (**Figure 2A**). Principal component analysis of RNA samples showed clear clustering of biological replicates (**Figure S3A**). The first principal component (PC1) separated samples by tissue and accounted for 70% of total expression variation, while PC2 separated samples primarily by species phylogeny and explained only 5% of variation (**Figure S3A**). Leaf and root expression patterns clustered first by tissue, then by species (**Figure S3A**). Leaf samples displayed greater variability than root samples (**Figure S3A**), and on PC2, species separated into two phylogenetic clusters: the Phaseoloids (*G. max* and *P. vulgaris*) plus *M. truncatula*, and the remaining Hologalegina species (*C. arietinum*, *L. culinaris*, *P. sativum*, and *L. sativus*) (**Figure S3A**).

**Figure 2.**
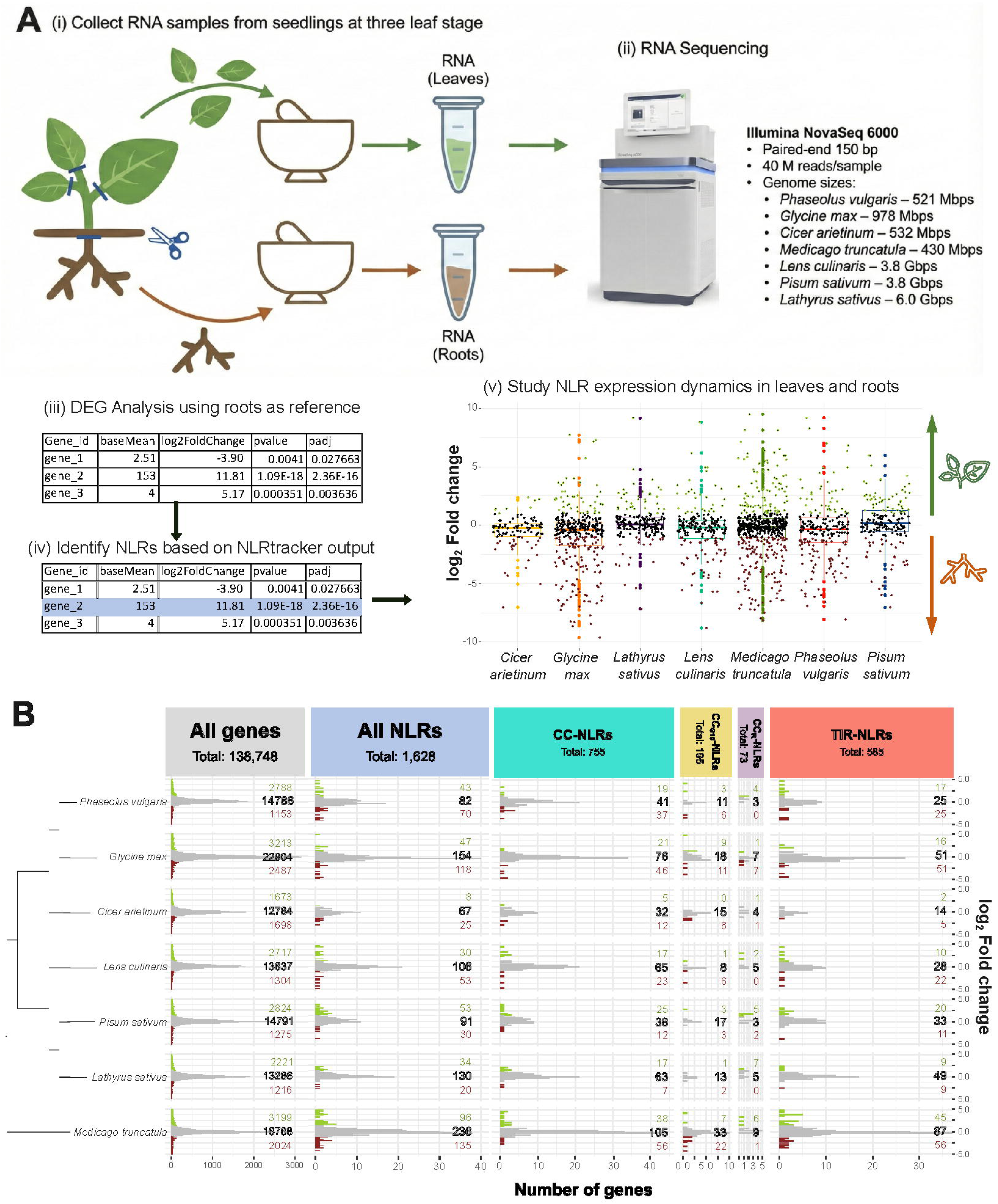
Tissue-specific expression patterns in legume transcriptomes. (**A**) Experimental design showing tissue collection from seedlings at the three-leaf developmental stage for RNA-seq analysis using Illumina NovaSeq 6000 (150 bp paired-end, 40M reads per sample), followed by DEG analysis, NLR identification, and analysis of NLR expression dynamics. (**B**) Tissue expression profiles across seven legume species. Horizontal histograms show gene frequency (x-axis) versus log_2_ fold change (y-axis). Positive values indicate leaf-predominant expression (green), negative values indicate root-predominant expression (brown). Absolute gene numbers are shown on the right.

Clustered heatmap analysis further supported this tissue-dependent pattern, with samples clustering primarily in two tissue groups, with closely related species forming distinct clusters within each group (**Figure S3B**). Differences between species were sufficiently large for the biological replicates from each species to cluster independently (**Figure S3B**). Of 3,782 BUSCO orthologues present across all seven species, 3,160 (84%) were expressed in both tissues. Notably, 1,075 of these genes demonstrated high average expression levels, exceeding 250 TMM.

Among the genes with highest leaf enrichment, we found *P. sativum*’s chloroplastic ribulose bisphosphate carboxylase/oxygenase activase (RuBisCO activase, LOC127083843, logFC = 15.06), *M. truncatula*’s chloroplastic carbonic anhydrase (MtrunA17_Chr6g0450641, logFC = 14.78), *C. arietinum*’s ribulose-bisphosphate carboxylase (RuBP carboxylase, Ca_16979, logFC = 14.54), *L. culinaris*’ chloroplastic RuBisCO activase (Lcu.2RBY.3g039110, logFC = 14.34), *L. sativus*’ chloroplastic RuBisCO activase (g19754, logFC = 14.13), *P. vulgaris*’ RuBP carboxylase (Phvul.004G064800, logFC = 13.33), and *G. max*’s light-harvesting complex II chlorophyll a/b binding protein 3 (LHCB3, Glyma.13G282000, logFC = 12.19).

On the other hand, the genes with the highest root enrichment were a *P. sativum* leghemoglobin (LOC127121418, logFC = -14.14), a *M. truncatula* nodulin-25 (MtrunA17_Chr3g0100391, logFC = -14.14), a *P. vulgaris* linoleate 9S-lipoxygenase-like gene (Phvul.005G156800, logFC = -12.97), the *L. sativus* gene g29944 (no predicted gene function or domains, logFC = -12.61), the highly root-expressed (according to Phytozome data; Goodstein *et al*., 2012) *G. max* gene Glyma.09G092700 (logFC = -12.12), a *L. culinaris* disease resistance response protein Pi49-like (Lcu.2RBY.2g016700, logFC = - 10.86), and a *C. arietinum* chitinase (Ca_18462, logFC = -9.95).

The top differentially expressed genes in leaves and roots are well-known tissue-specific markers, supporting the robustness of the transcriptomic data.

Tissue identity drives substantially greater expression variation than species differences, providing a framework for interpreting tissue expression patterns of genes.

### NLRs exhibit enhanced tissue-specificity with predominant root expression

NLR tissue-specificity patterns may differ from genome-wide patterns due to their specialized immune function. To investigate NLR tissue-specificity compared to other gene families, we analyzed differential expression patterns across the seven legume species.

Among 138,748 expressed genes across the seven species, 21.47% exhibited differential expression between tissues (**Figure 2B** – All genes), with leaf expression favored 1.68 to 1 (**Table S8**). Transcriptomic NLR analysis revealed that 1,628 of 1,858 predicted NLRs were expressed in at least one tissue, with 1,336 sufficiently expressed transcripts for DEG analysis (71.9% expression rate). Remarkably, 57.04% of expressed NLRs showed differential expression between tissues (**Figure 2B** – All NLRs), with root expression favored 1.45 to 1 (**Table S8**). Legume NLRs displayed 2.66-fold greater tissue-dependent expression than all gene families combined and reversed the general leaf-biased tendency, showing predominant root expression instead.

Root-specific NLR expression exceeded leaf-specific expression, particularly in *M. truncatula*, *G. max*, and *P. vulgaris* (**Figure 2B, S4**). However, both *L. sativus* and *P. sativum* showed a tendency towards NLR leaf-specific expression instead (**Figure 2B, S4**). On average, 21% of expressed NLRs were root-specific, with *M. truncatula* showing the highest proportion (30%) and *L. sativus* the lowest (8%) (**Figure 3B**). Leaf-specifically expressed NLRs averaged 11.5% across species, with *P. sativum* displaying the highest proportion (16%) and *C. arietinum* the lowest (2%) (**Figure 3B**). Most expressed NLRs (50-83%) were detected in both tissues across all species (**Figure 3B**).

**Figure 3.**
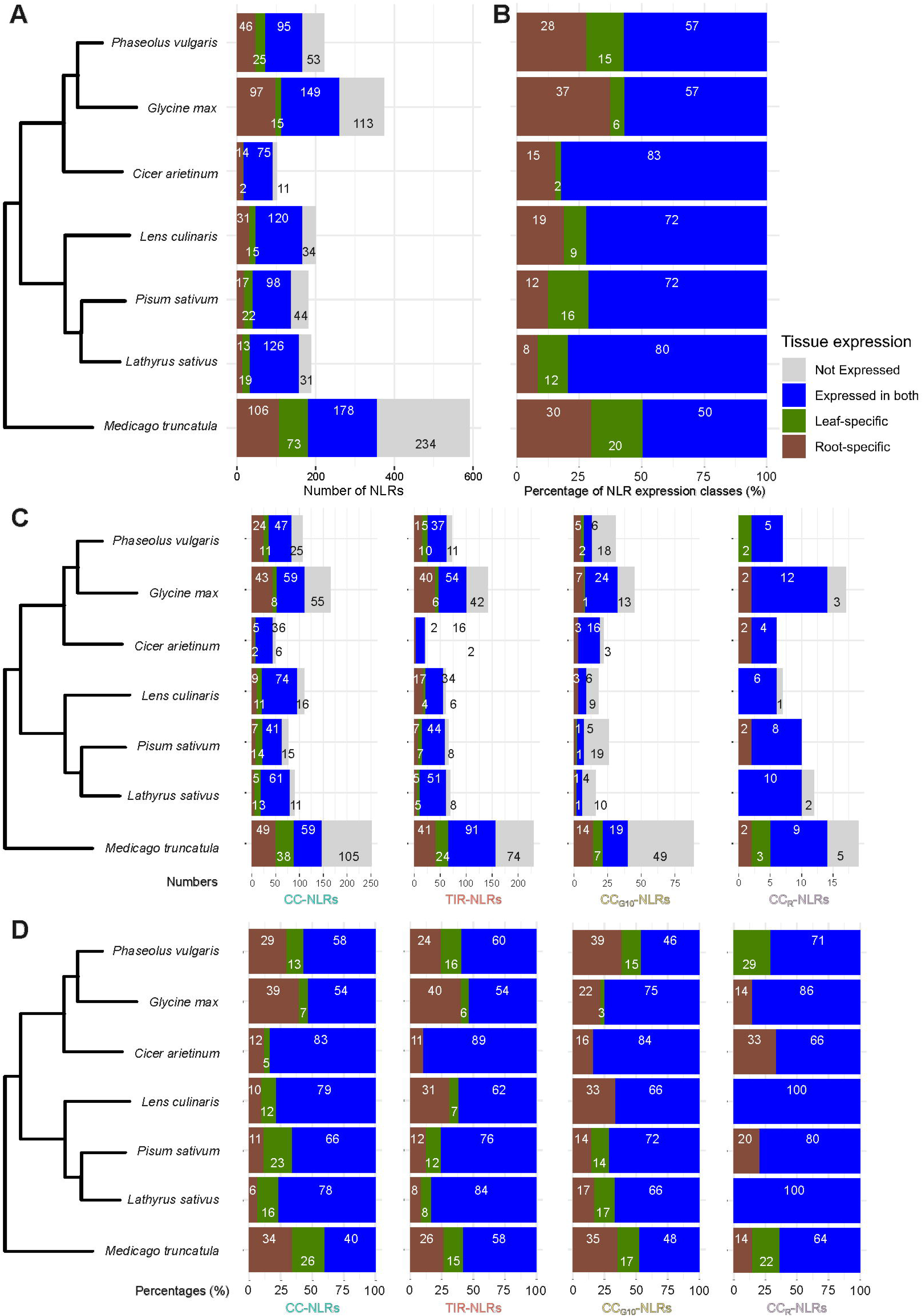
Stacked bar charts showing NLR expression patterns across tissue and species. (**A**) Tissue expression patterns of NLRs in absolute numbers. (**B**) Percentages of tissue expression patterns across expressed NLRs. (**C**) Stacked bar charts showing tissue expression patterns of each NLR class (CC-NLRs, TIR-NLRs, CC_G10_-NLRs, and CC_R_-NLRs). (**D**) Percentages of tissue expression patterns across the classes of expressed NLRs (CC-NLRs, TIR-NLRs, CC_G10_-NLRs, and CC_R_-NLRs). All figures share the same legend: Grey bars indicate non-expressed NLRs; blue bars show NLRs expressed in either tissue (non-tissue-specific); brown bars show root-specific NLRs; and green bars show leaf-specific NLRs.

NLR class-specific analysis revealed distinct tissue predominance among expressed NLRs (**Table S8**). CC_R_-NLRs showed leaf bias with 40% (27/67) expressing predominantly in leaves versus 19% (13/67) in roots, particularly in *L. sativus*, *P. sativum*, and *P. vulgaris* (**Table S8**). TNPs demonstrated strong root-bias with 86.7% (13/15) showing root-specific expression (**Table S7**). CC_G10_-NLRs favored roots with 45.2% (57/126) root-specificity versus 21.4% (27/126) leaf-specificity (**Table S8**). CC-NLRs and TIR-NLRs showed similar patterns with ∼25-26% leaf-specificity (158/616 and 130/512) and ∼32-35% root-specificity (198/616 and 181/512) (**Figure 2B**). Surprisingly, TIR-NLRs and their required CC_R_-NLR helper-NLRs (NRG1 and ADR1 subfamilies) showed opposing tissue predominance. While most TIR-NLRs were root-specific, their helper-NLRs were predominantly expressed in leaves.

*Medicago truncatula* and *C. arietinum* were on opposite sides of the NLRome size spectrum. *Cicer arietinum*, the species with the smallest NLRome, showed the lowest proportion of overall non-expressed NLRs (11%) and tissue-specific NLRs (17%; **Figure S4E**). On the other hand, *M. truncatula*, the species with the largest NLRome, exhibited the highest proportions of non-expressed NLRs (40%) and tissue-specificity (50%; **Figure S4E**).

NLRs exhibit enhanced tissue-specificity compared to other gene families and show predominant root expression that contrasts with genome-wide leaf bias patterns.

### Evolutionary conservation of tissue expression in NLR subclasses

Tissue-enriched NLR subclasses can reveal evolutionary conservation and identify potential functional groups. To identify conserved tissue-enriched NLR subclasses, we examined prominent phylogenetic subclasses spanning NLRs from multiple species and assessed whether these groups exhibited consistent leaf- or root-enriched expression patterns (**Figure 4**).

**Figure 4.**
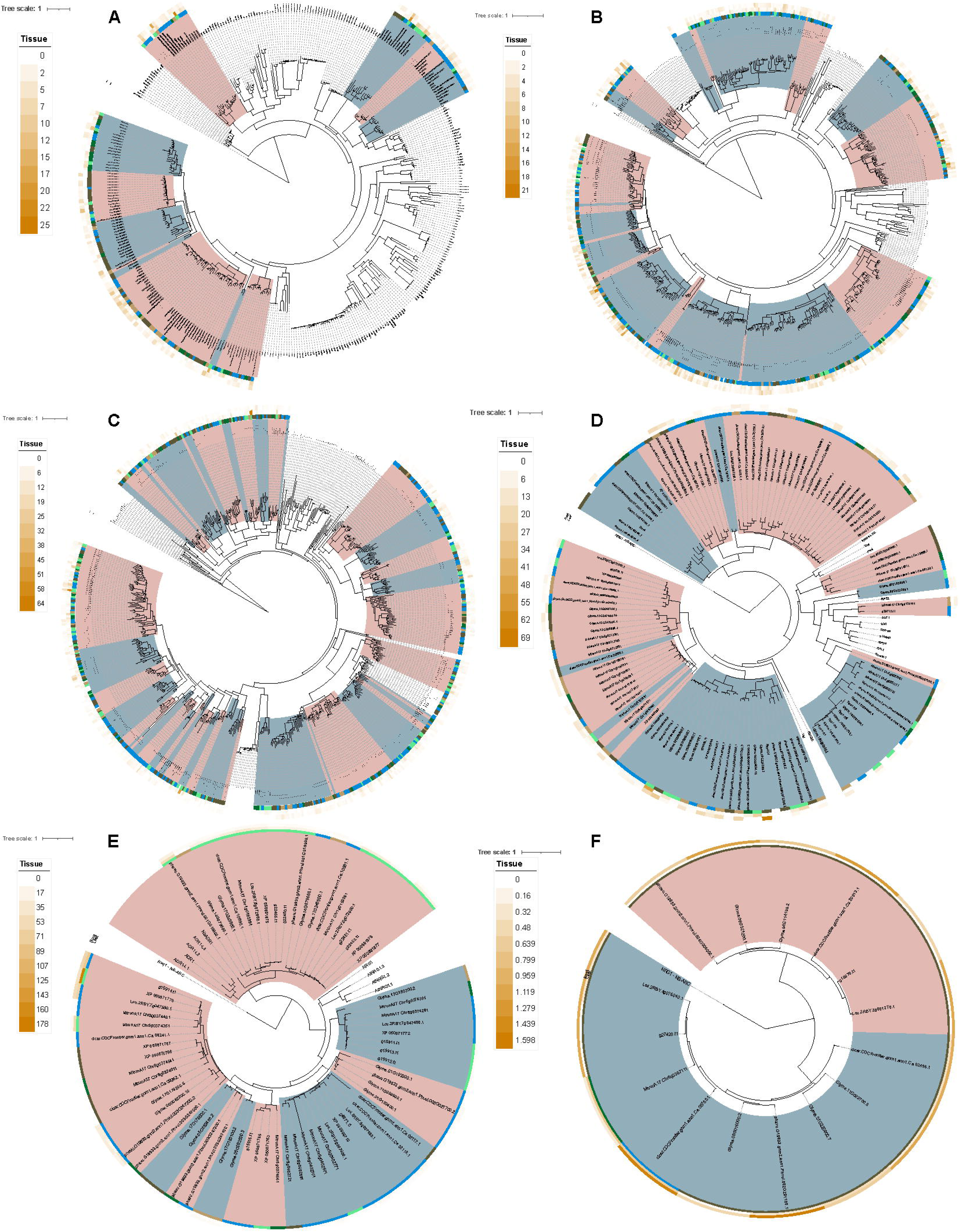
Phylogenetic trees of cross-species NLR subclasses including RefPlantNLR entries. (**A**) Phylogenetic tree of 164 CC-clade-1 NLRs divided into 12 subclasses. (**B**) Phylogenetic tree of 437 CC-clade-2 NLRs divided into 21 subclasses. (**C**) Phylogenetic tree of 506 TIR-NLRs divided into 51 subclasses. (**D**) Phylogenetic tree of 112 CC_G10_-NLRs divided into 17 subclasses. (**E**) Phylogenetic tree of 67 CC_R_-NLRs divided into an ADR1 subclass and eight NRG1 subclasses. (**F**) Phylogenetic tree of 15 TNPs divided into 2 subclasses. The clade colors separate NLR subclasses within each NLR class phylogenetic tree. Figure generated using iTOL v6 (Letunic and Bork, 2024).

Our analysis revealed 113 NLR subclasses (**Table S10**). Among these, twelve had strong cross-species enrichment signatures, spanning CC_R_-, CC-, and TIR-NLRs. Among helper-NLRs, ADR1-01 displayed conserved leaf specialization, comprising 19 NLRs across all seven species, of which 15 were leaf-enriched. Multiple CC-NLR subclasses also showed consistent expression bias. CC-clade1-07 was strongly root-enriched (25 root-enriched of 42 NLRs across all seven species). CC-clade1-10 contained 16 NLRs across four species, with 12 showing root enrichment, whereas CC-clade1-11 included 17 NLRs across two species, of which 11 were root-enriched. In contrast, CC-clade1-12 represented a leaf-biased CC-NLR clade (13 leaf-enriched from 19 NLRs across four species). An additional CC-NLR group, CC-clade2-06, showed near-uniform root enrichment (6 of 7 NLRs across four species).

TIR-NLR subclasses likewise exhibited conserved tissue patterns (**Table S10**). Several TIR-NLR subclasses showed predominant root enrichment: TIR-02 (6 root-enriched of 10 NLRs across five species), TIR-11 (14 root-enriched of 22 NLRs across all seven species), TIR-38 (3 of 5 NLRs across five species), and TIR-43, a large root-biased lineage composed of 32 NLRs across two species, with 21 showing predominant root enrichment. Leaf-biased TIR-NLR subclasses were also present, including TIR-05 (with 3 of 4 leaf-enriched NLRs across four species) and TIR-19 (8 leaf-enriched of 9 NLRs across six species).

Overall, the recurrence of tissue-enriched expression within phylogenetically close NLR subclasses indicates that leaf- or root-associated specialization is not randomly distributed but can reflect evolutionary patterns shared across legume species. These conserved subclasses provide a focused set of candidate lineages for downstream functional or evolutionary analyses.

### Tissue-specific NLRs cluster by species phylogeny and NLR class

Tissue-specific gene expression patterns may be evolutionarily conserved or may evolve independently in different lineages. To further investigate whether tissue-specific NLR expression patterns are phylogenetically conserved, we performed phylogenetic analysis of tissue-specific NLRs across seven species.

Phylogenetic analysis of leaf-specific (**Figure 5A**) and root-specific (**Figure 5B**) NLRs revealed clustering patterns driven by NLR class and interspecific phylogenetic distance. NLRs from Hologalegina species (*M. truncatula*, *C. arietinum*, *L. culinaris*, *P. sativum*, and *L. sativus*) cluster together, while Phaseoloids (*P. vulgaris* and *G. max*) form separate clusters (**Figure 5**). Root-specific NLRs showed markedly higher expression levels than leaf-specific NLRs (**Figure 5**). Many clusters lacked representatives from some species, indicating either species-specific expansions, gene loss, or incomplete genomic resources. Root-specific CC_R_-NLRs belonged to the NRG1 subfamily, with homologues identified in four species (**Table S9**). TNPs showed root-specific expression in five species, whereas *M. truncatula* was the only species with a leaf-specific TNP (**Figure 5A, Table S9**).

**Figure 5.**
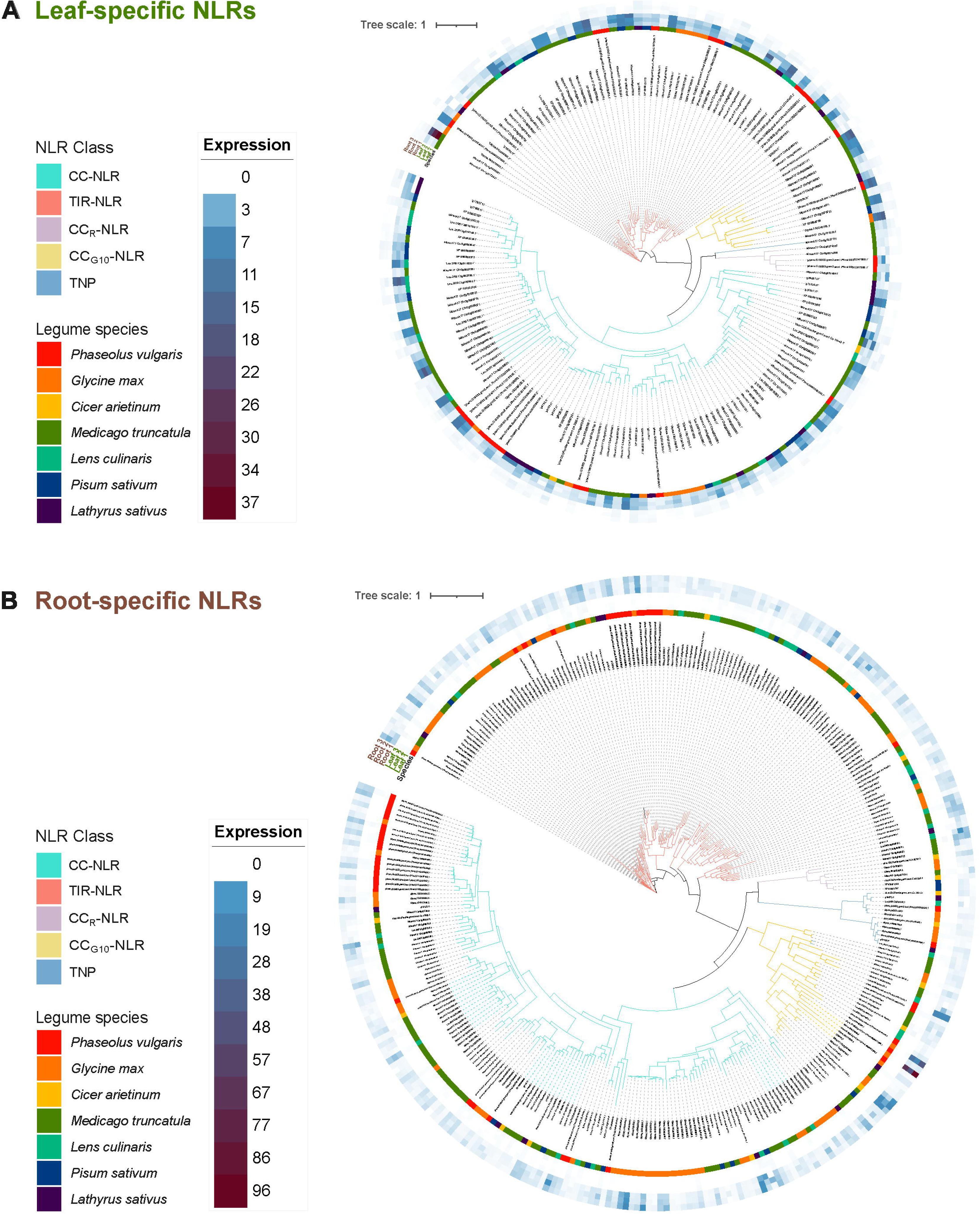
Phylogenetic relationships and expression patterns of tissue-specific NLRs. Expression values in trimmed mean of M-values (TMM) across three biological replicates for (**A**) leaf-specific and (**B**) root-specific NLRs. Species colors: *P. vulgaris* (red), *G. max* (orange), *C. arietinum* (yellow), *M. truncatula* (green), *L. culinaris* (teal), *P. sativum* (blue), *L. sativus* (purple). NLR class colors are represented in the color of tree branches: turquoise (CC-NLRs), salmon (TIR-NLRs), lilac (CC_R_-NLRs), yellow (CC_G10_-NLRs), and blue (TNPs). Expression scale: white (0 TMM), blue (10 TMM), dark red (respective maximum values). Figure generated using iTOL v6 (Letunic and Bork, 2024).

## DISCUSSION

In this study, we demonstrate that the expression of NLR immune receptors in legumes can be highly compartmentalized by organ tissue and provide a valuable community resource to support interspecific comparative studies. Legume NLRs exhibit over twice the level of tissue-specific expression compared to the genomic average, generally with a skew towards root tissues. This finding contrasts with the overall leaf-biased expression of most other genes, suggesting that distinct selective pressures in the rhizosphere may have shaped the evolution of the legume NLR immune system. This root-centric defensive investment in legumes aligns with observations in *Lotus japonicus* and monocots (Tarr and Alexander, 2009; Munch *et al*., 2018), but is opposite to the shoot-skewed NLR expression reported in Brassicaceae, a group that lost the capacity for root endosymbiosis with mycorrhizal fungi (Lai and Eulgem, 2018; Munch *et al*., 2018), highlighting potential divergence in immune compartmentalization between these lineages. The tissue deployment of distinct NLR classes, such as the predominant expression of CC_R_-NLRs in leaves and TNPs in roots, further points to the spatial organization of plant defense.

The root bias in NLR expression may reflect an evolutionary adaptation to the complex and microbe-rich soil environment (Munch *et al*., 2018). Roots face constant exposure to a diverse array of soil-borne pathogens and microorganisms, requiring a large and varied immune receptor repertoire. Furthermore, the unique symbiotic relationship between legumes and nitrogen-fixing rhizobia imposes an additional layer of complexity (Zhao *et al*., 2021). The immune system allows plants to differentiate between pathogens and beneficial microbes, and NLRs are known to act as regulators of host-rhizobia specificity (Yang *et al*., 2010a, 2010b; Kisseleva, 2014; Quilbé *et al*., 2021). The abundance of root-expressed NLRs may therefore provide a molecular toolkit not only for pathogen defense but also for modulating symbiosis. The identification of root-specific NLRs clustering with receptors known to confer resistance to soil-borne diseases, such as Rps1-k (*Phytophthora sojae* root rot), supports their role in subterranean defense and provides promising candidates for future functional validation(Gao *et al*., 2005). It is also noteworthy that the present NLR expression patterns were observed in plants grown in non-aseptic conditions, with roots containing nodules, and were therefore shaped by responses to microbes present in the phyllo- and rhizosphere.

While root-biased expression is the dominant trend, our findings reveal important exceptions that underscore the influence of lineage-specific evolution. The closely related species *P. sativum* and *L. sativus* displayed predominantly leaf-specific NLR expression, opposite to the other five species analyzed. Since nodulation was similar across all species at the time of tissue collection, this distinct pattern is unlikely to reflect differences in symbiotic status and instead suggests that tissue-specific expression patterns can be conserved within lineages. The distinct pattern of these Fabeae tribe species (Schaefer *et al*., 2012; Santos and Leitão, 2023) may reflect adaptation to historical or ongoing pressure from phyllosphere-specific pathogens (Rubiales *et al*., 2015). It could also have resulted from a depletion of NLRs expressed in roots to allow symbiosis with a wider diversity of rhizobia (Benezech *et al*., 2020).

Additionally, we observed a striking discordance in tissue predominance between TIR-NLRs, which were largely root-specific, and their canonical CC_R_-NLR helper-NLRs (*NRG1/ADR1* families), which were predominantly expressed in leaves and tend to be constitutively expressed. This pattern suggests two possible scenarios: distinct pools of helper-NLRs may operate in different tissues, with a subset sufficient to support TIR-NLR-mediated immunity in each location; alternatively, a subset of broadly expressed helper-NLRs may be sufficient to support TIR-NLR function across the entire plant, representing an efficient allocation of defensive resources (Jubic *et al*., 2019). The high expression values observed for CC_R_-NLRs relative to other NLR classes are consistent with their role as central, conserved hubs in immunity that must be readily available for signal transduction (Castel *et al*., 2019; Jubic *et al*., 2019; Sun *et al*., 2021; Contreras *et al*., 2023).

Our analysis of NLRome size in relation to expression patterns reveals a potential evolutionary trade-off. Species with the largest NLRomes, such as *M. truncatula*, also exhibited the highest proportion of both non-expressed and tissue-specific NLRs. Conversely, *C. arietinum*, with one of the smallest NLRomes, showed broad expression across tissues with minimal tissue-specificity. This suggests that NLRome expansion may impose a fitness cost (Richard *et al*., 2018) that results in an increased compartmentalization. Consequently, larger NLRomes appear to display transcriptional silencing or strict tissue-specific regulation, whereas smaller, more streamlined NLRomes may rely on widespread expression to provide protection (Richard *et al*., 2018). This aligns with the “birth-and-death” model of NLR evolution, where expansions are coupled with functional specialization and regulatory refinement (Michelmore and Meyers, 1998; Wu *et al*., 2017). This pattern may also be influenced by agricultural history; the smaller NLRome in a domesticated crop that suffered major genetic bottlenecks, like *C. arietinum* (Abbo *et al*., 2003; Raina *et al*., 2019), could be a consequence of “NLRome erosion”, a process where intense selection for yield and other traits during breeding leads to the inadvertent loss of disease resistance genes that are more abundant in wild relatives (Meyer *et al*., 2012; Flint-Garcia, 2013; Zheng *et al*., 2016; Ma *et al*., 2019; Gasparini *et al*., 2021; Bayer *et al*., 2022; Asif *et al*., 2023).

Constitutive NLR expression across major plant organs and tissues may reflect an adaptive strategy for this fast-evolving gene superfamily (Zheng *et al*., 2016; Rani *et al*., 2023), as expression in multiple tissues could maximize the potential benefits of any gain-of-function mutations. Furthermore, the legume family experienced several whole genome duplications across its evolution and domestication, accompanied by purifying selection forces (Zheng *et al*., 2016; Zhao *et al*., 2021). The genome dynamics in the legume family could explain the expansion and contraction of NLR repertoires across species (Zheng *et al*., 2016; Qureshi *et al*., 2023). Despite the variable quality of the legume genomic resources, we observed a significant positive correlation between proteome size and NLR number, contrasting with similar studies on legume NLRomes (Qureshi *et al*., 2023; Negi *et al*., 2024). The number of NLRs per species identified in the present study is substantially less (often around a third) than those reported in Negi *et al*. (2024), likely because of the strict filtering we employed to avoid double counting splicing variants and masking truncated NLRs that lack canonical domains. Nevertheless, the proportion of protein-coding loci encoding NLRs aligns with those obtained in Ngou *et al*. (2022).

This study establishes a framework for understanding the spatial deployment of the legume immune system across organs and tissues, while also serving as a community resource and springboard for future investigation, including comparative interspecific studies on NLR expression and breeding for tissue-specific disease resistance. The variable quality of some publicly available legume genomes may have limited the complete identification of all NLRs, and future work with improved assemblies will refine these NLR repertoires. Our analysis was conducted on unchallenged seedlings at a single developmental stage. Transcriptomic profiling across different life stages and in response to pathogen infection (like the studies in Fick *et al*., 2022, and Maravilha *et al*., 2025) is needed to capture the dynamic nature of NLR expression. Finally, the NLR subclasses and clades identified here, especially the ones showing tissue expression conservation across species, are prime targets for functional characterization. Validating their roles in disease resistance will be crucial for translating these findings into applied breeding programs, enabling the development of durable, bioengineered resistance targeted to the tissues where pathogenic challenges are anticipated.

## Acknowledgements

The authors thank the following individuals for generously providing seeds: Dr. Sanu Arora (John Innes Centre, Norwich, UK) for *Pisum sativum* Caméor; Dr. Myriam Charpentier (John Innes Centre, Norwich, UK) for *Medicago truncatula* A17; Dr. Sara Dorhmi (The Sainsbury Laboratory, Norwich, UK) for *Glycine max* Williams 82; and Dr. Bunyamin Taran (University of Saskatchewan, Western Canada) for *Cicer arietinum* CDC Frontier. We also thank Dr. Kirstin E. Bett (University of Saskatchewan, Western Canada) for generously providing the most recent genome assembly and seeds for *Lens culinaris* CDC Redberry. Seeds for *Phaseolus vulgaris* G19833 were provided by the Genetic Resource Unit from the International Center for Tropical Agriculture (CIAT). We thank Dr. Daniel Lüdke and AmirAli Toghani for their valuable input on RNA-Seq data analysis and visualization. We also acknowledge Dr. Yu Sugihara and Dr. Joe Win for assistance with NLRtracker analysis.

## Author contributions

**R.M.M.:** Conceptualization, Methodology, Formal Analysis, Investigation, Data Curation, Writing – Original Draft, Visualization. **C.S.:** Conceptualization, Supervision, Writing – Review & Editing. **H.P.:** Investigation, Writing – Review & Editing. **M.C.V.P.:** Conceptualization, Supervision, Writing – Review & Editing. **S.K.:** Conceptualization, Supervision, Project Administration, Funding Acquisition, Writing – Review & Editing. **J.K.:** Methodology, Software, Formal Analysis, Supervision, Writing – Review & Editing.

## Data availability

The raw RNA-seq data and corresponding metadata generated in this study have been deposited in the ArrayExpress database at EMBL-EBI under accession number E-MTAB-15524. All NLR annotation files, data, and related analysis outputs are available at Zenodo [10.5281/zenodo.16852232] (Maravilha Marques *et al*., 2026). All other data supporting the findings of this study are available within the manuscript and its supplementary information files.

## SUPPORTING INFORMATION

**Supplemental Figure 1. Phylogenetic relationships and genome quality assessment of species used in this study. (A)** Phylogenetic tree based on three chloroplast genes (*atpA*, *matK*, and *rbcL*) confirms expected evolutionary relationships among 23 legume species and four outgroups. **(B)** Genome assembly contiguity varies across species. Contig N50 values are shown on a log scale, with the dashed line indicating the 1 Mbp threshold for a highly contiguous assembly. (**C**) Predicted proteome completeness shows significant variation. BUSCO scores were calculated using the fabales_odb10 (legumes) or embryophyta_odb10 (outgroups) datasets. The dashed line indicates the 95% completeness threshold. Bar colors represent BUSCO categories: complete and duplicated (salmon), complete and single-copy (orange), fragmented (light blue), and missing (dark blue).

**Supplemental Figure 2. NLR gene counts are distributed unevenly across canonical NLR classes.** Boxplots show the distribution of gene counts for each NLR class across the 28 analyzed legume species. The central line indicates the median, box limits represent the upper and lower quartiles, and whiskers extend to 1.5 times the interquartile range. CC-NLRs and TIR-NLRs are the most abundant and widely distributed classes.

**Supplemental Figure 3. Tissue identity is the primary driver of gene expression variation.** Analysis of BUSCO gene expression across seven legume species in leaf (L) and root (R) tissues for *Cicer arietinum* (Car), *Glycine max* (Gma), *Lathyrus sativus* (Lsa), *Lens culinaris* (Lcu), *Medicago truncatula* (Mtr), *Phaseolus vulgaris* (Pvu), and *Pisum sativum* (Psa). **(A)** Principal component analysis (PCA) of BUSCO gene expression. Samples cluster primarily by tissue (PC1, 70% variance) and secondarily by species phylogeny (PC2, 5% variance). **(B)** Hierarchical clustering of BUSCO gene expression. Samples (columns) cluster first into distinct tissue groups and then by species. Genes (rows) are clustered by expression similarity. The heatmap is colored by log-transformed TMM values.

**Supplemental Figure 4. Species-specific analysis reveals patterns of NLR expression and tissue predominance.** Circular phylogenies displaying NLR expression for each of the seven species: **(A)** *Cicer arietinum*, **(B)** *Glycine max*, **(C)** *Lens culinaris*, **(D)** *Lathyrus sativus*, **(E)** *Medicago truncatula*, **(F)** *Pisum sativum*, and **(G)** *Phaseolus vulgaris*. For each plot, the concentric rings display, from innermost to outermost: **(i)** NLR class (CC-NLR: turquoise; TIR-NLR: salmon; CC_G10_-NLR: yellow; CC_R_-NLR: lilac; TNP: blue), **(ii)** TMM expression values across three leaf and three root biological replicates, and **(iii)** tissue-specificity classification (expressed in both tissues: blue; leaf-specific: green; root-specific: brown; not expressed: black).

**Supplemental Table 1.** Quality parameters and origin of all genomes used in the study.

**Supplemental Table 2.** BUSCO gene (fabales_odb) counts across samples.

**Supplemental Table 3.** NLRs with RefPlantNLR subdivided into the NLR classes CC-NLR, TIR-NLR, CC_G10_-NLR, and CC_R_-NLR.

**Supplemental Table 4.** Locus and NLR class for each NLR across species.

**Supplemental Table 5.** Summary table of NLR classes across species.

**Supplemental Table 6.** Expression data for all predicted NLRs in TMM values.

**Supplemental Table 7.** Summary table of expression data for all predicted NLRs.

**Supplemental Table 8.** Summary table of expression patterns per species for classes: all genes, all NLRs, CC-NLRs, TIR-NLRs, CC_G10_-NLRs, and CC_R_-NLRs. Describes absolute and relative numbers within each class and species, and leaf to root odds. Green colored odds show increased leaf expression, while brown-colored odds show increased root expression.

**Supplemental Table 9.** NLR subclasses across species. #N/A - RefPlantNLR entry.

**Supplemental Table 10.** Tissue expression summary of cross-species NLR subclasses.

