## Supplemental Figures for "NLR immune receptors can exhibit tissue-specific expression patterns across legume species"

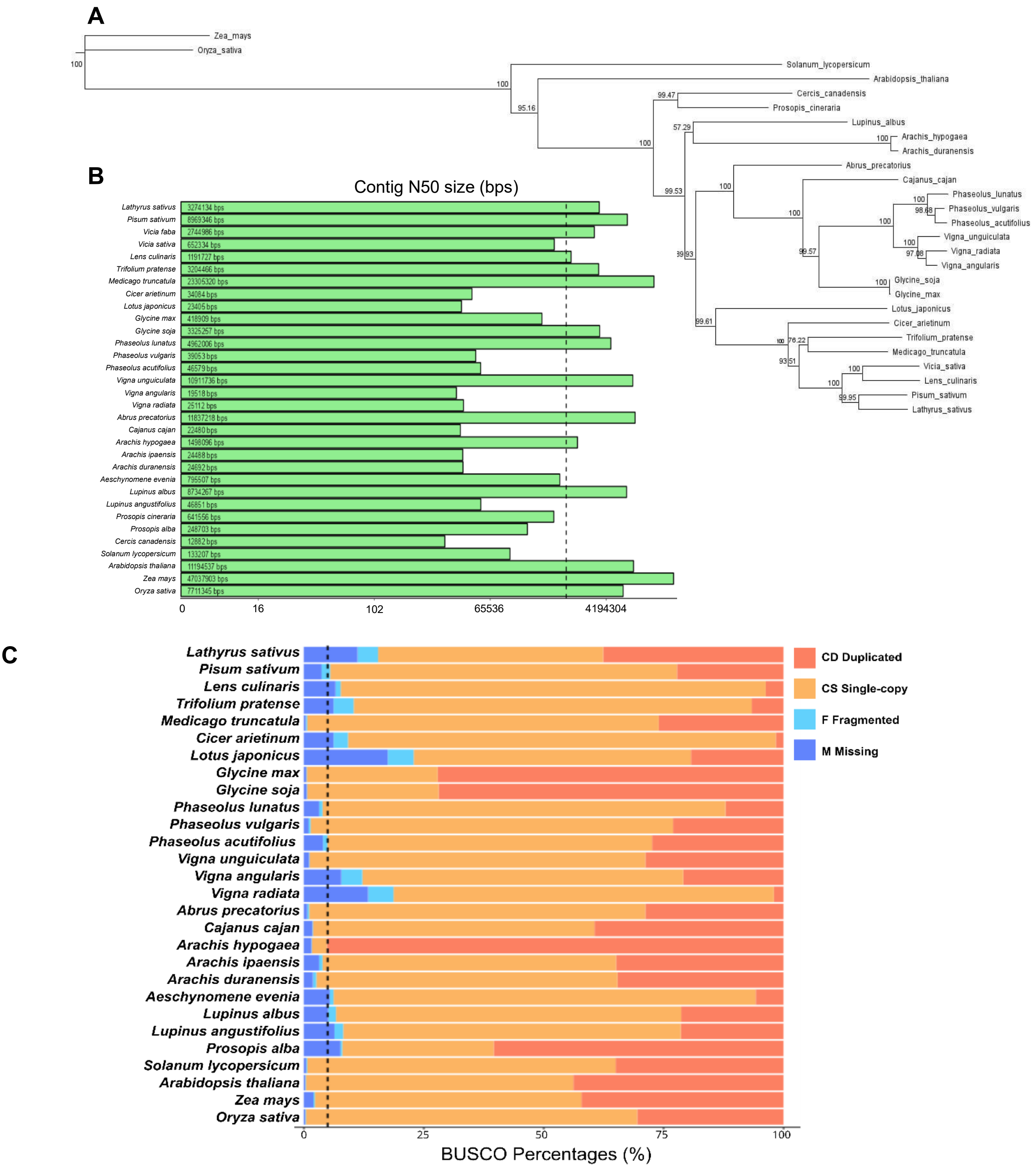


**Supplemental Figure 1. Phylogenetic relationships and genome quality assessment of species used in this study. (A)** Phylogenetic tree based on three chloroplast genes (*atpA*, *matK*, and *rbcL*) confirms expected evolutionary relationships among 27 legume species and four outgroups. **(B)** Genome assembly contiguity varies across species. Contig N50 values are shown on a log scale, with the dashed line indicating the 1 Mbp threshold for a highly contiguous assembly. (**C**) Predicted proteome completeness shows significant variation. BUSCO scores were calculated using the fabales_odb10 (legumes) or embryophyta_odb10 (outgroups) datasets. The dashed line indicates the 95% completeness threshold. Bar colors represent BUSCO categories: complete and duplicated (salmon), complete and single-copy (orange), fragmented (light blue), and missing (dark blue).

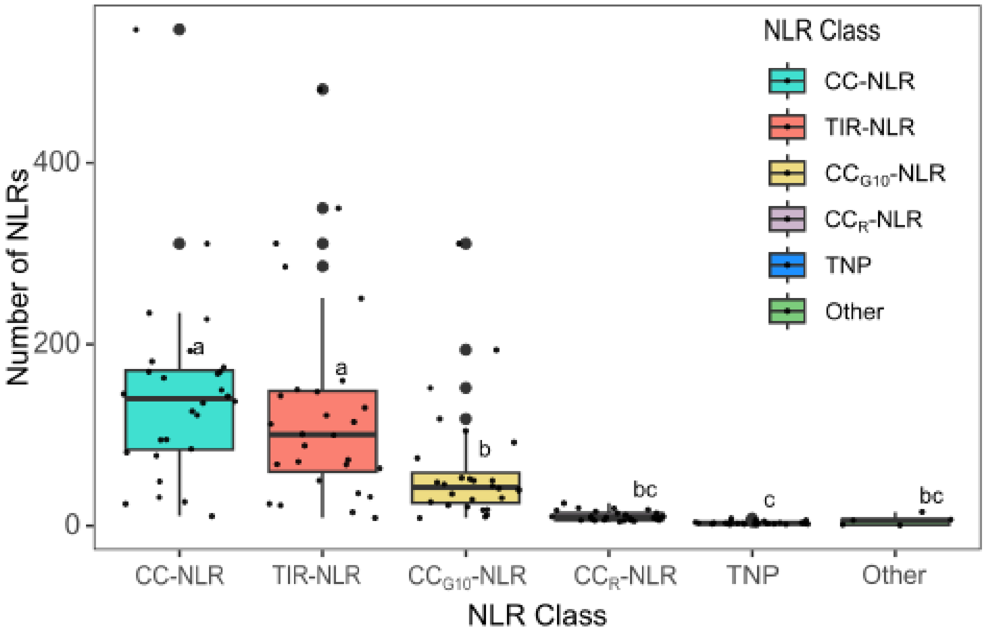


**Supplemental Figure 2. NLR gene counts are distributed unevenly across canonical NLR classes.** Boxplots show the distribution of gene counts for each NLR class across the 28 analyzed legume species. The central line indicates the median, box limits represent the upper and lower quartiles, and whiskers extend to 1.5 times the interquartile range. CC-NLRs and TIR-NLRs are the most abundant and widely distributed classes.

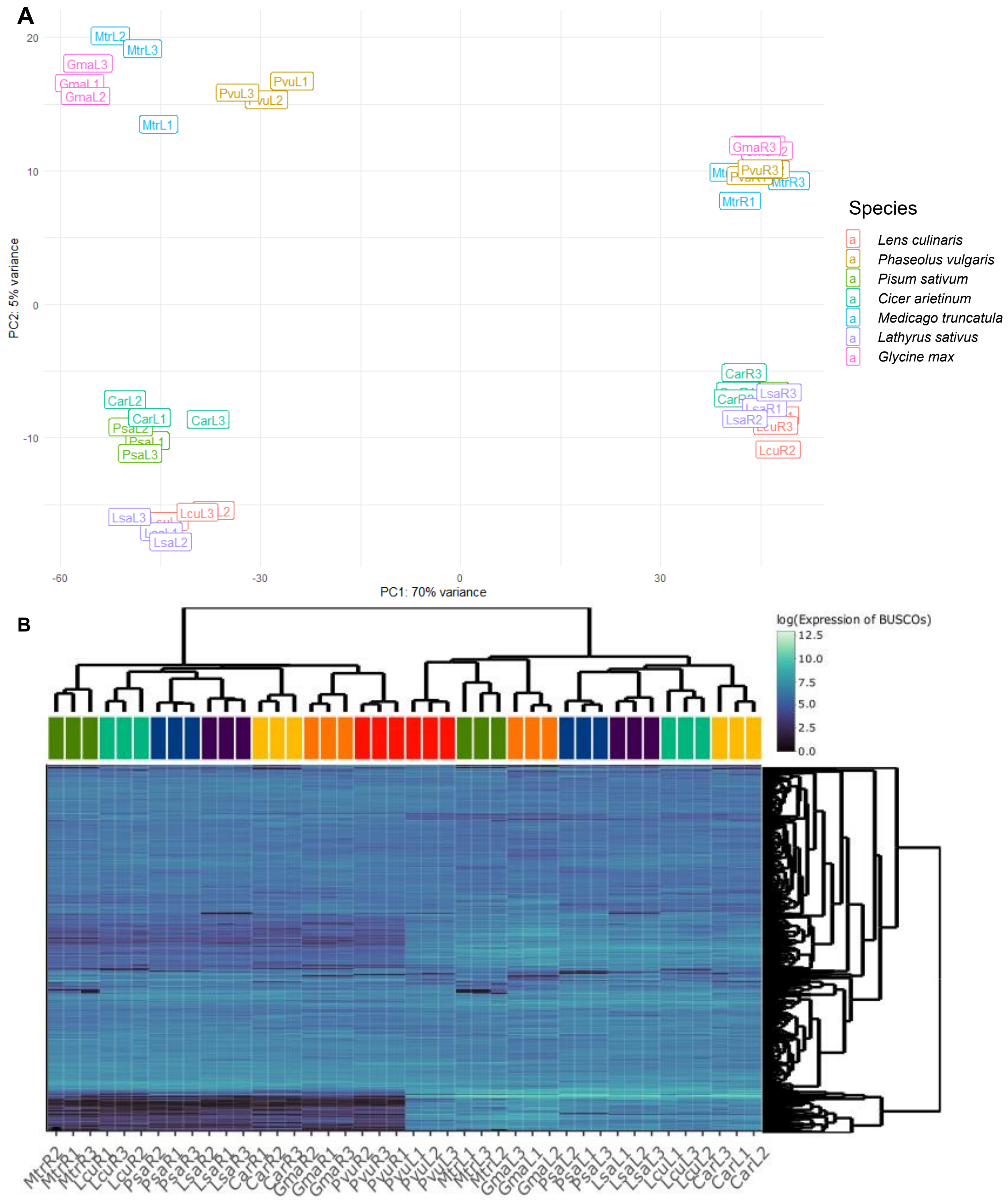


**Supplemental Figure 3. Tissue identity is the primary driver of gene expression variation.** Analysis of BUSCO gene expression across seven legume species in leaf (L) and root (R) tissues for *Cicer arietinum* (Car), *Glycine max* (Gma), *Lathyrus sativus* (Lsa), *Lens culinaris* (Lcu), *Medicago truncatula* (Mtr), *Phaseolus vulgaris* (Pvu), and *Pisum sativum* (Psa). **(A)** Principal component analysis (PCA) of BUSCO gene expression. Samples cluster primarily by tissue (PC1, 70% of variance) and secondarily by species phylogeny (PC2, 5% of variance). **(B)** Hierarchical clustering of BUSCO gene expression. Samples (columns) cluster first into distinct tissue groups and then by species. Genes (rows) are clustered by expression similarity. The heatmap is colored by log-transformed TMM values.

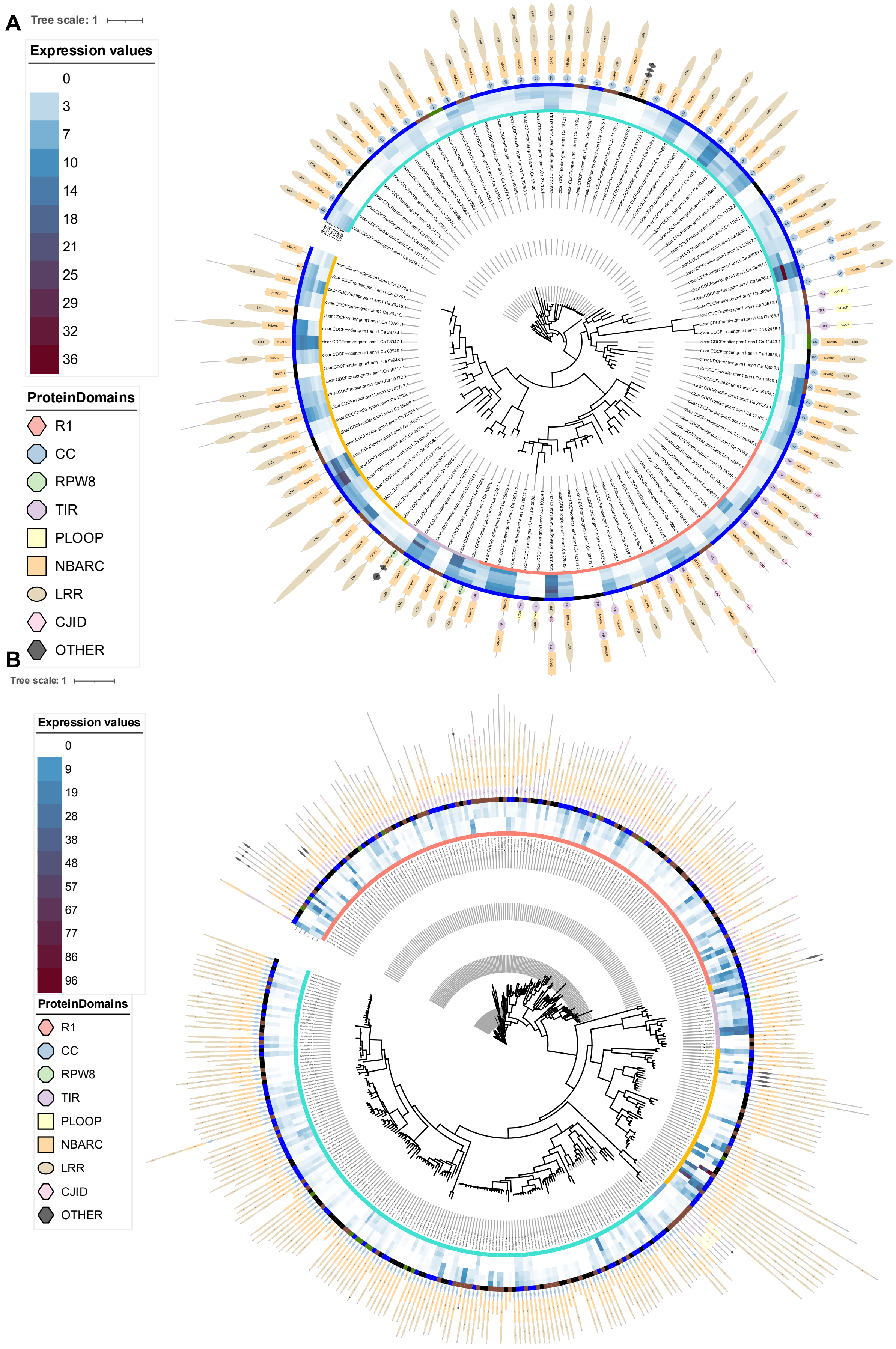
A
Tree scale: 1
Expression values
0
3
7
10
14
18
21
25
29
32
36
ProteinDomains
R1
CC
RPW8
TIR
PLOOP
NBARC
LRR
CJID
OTHER
B
Tree scale: 1
Expression values
0
9
19
28
38
48
57
67
77
86
96
ProteinDomains
R1
CC
RPW8
TIR
PLOOP
NBARC
LRR
CJID
OTHER

C

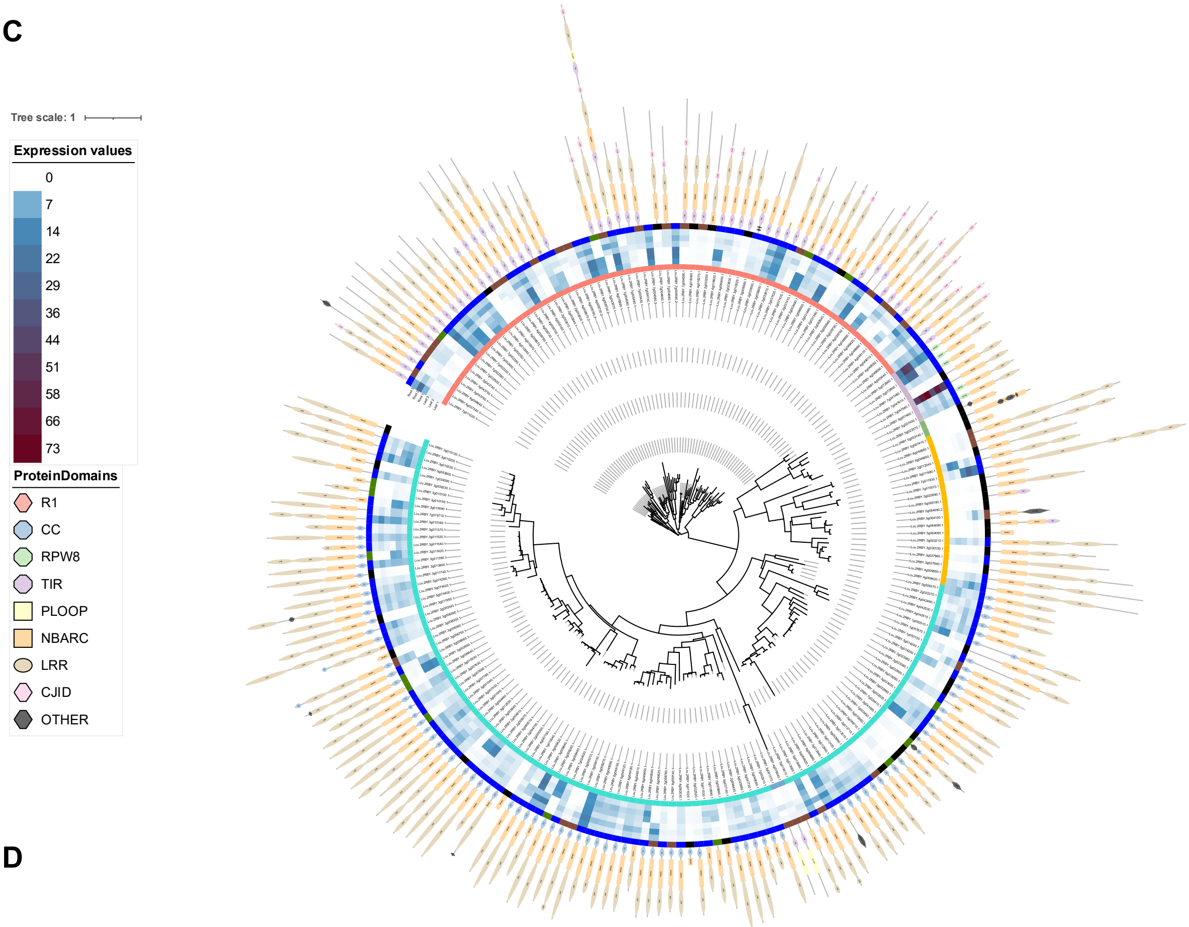

Tree scale: 1
Expression values
0
7
14
22
29
36
44
51
58
66
73
ProteinDomains
R1
CC
RPW8
TIR
PLOOP
NBARC
LRR
CJID
OTHER


D

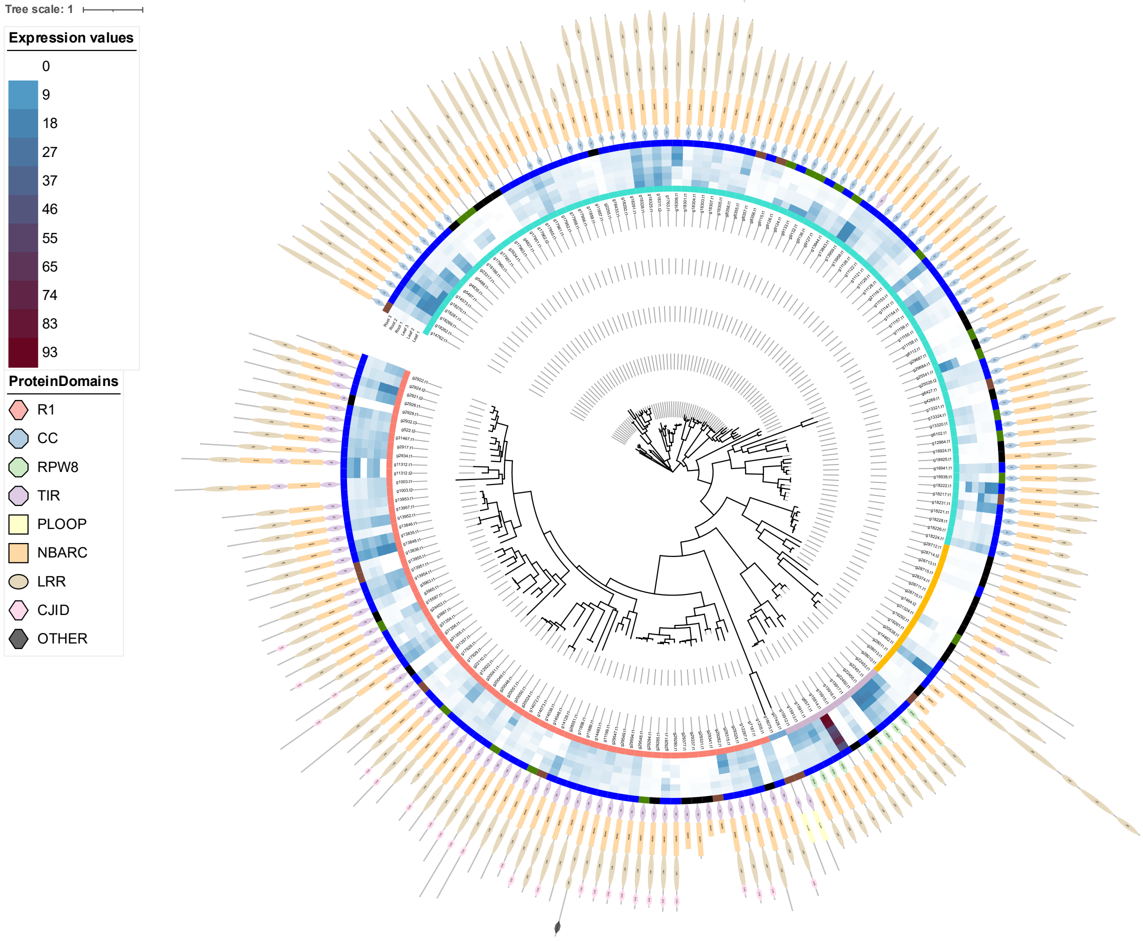

Tree scale: 1
Expression values
0
9
18
27
37
46
55
65
74
83
93
ProteinDomains
R1
CC
RPW8
TIR
PLOOP
NBARC
LRR
CJID
OTHER

E

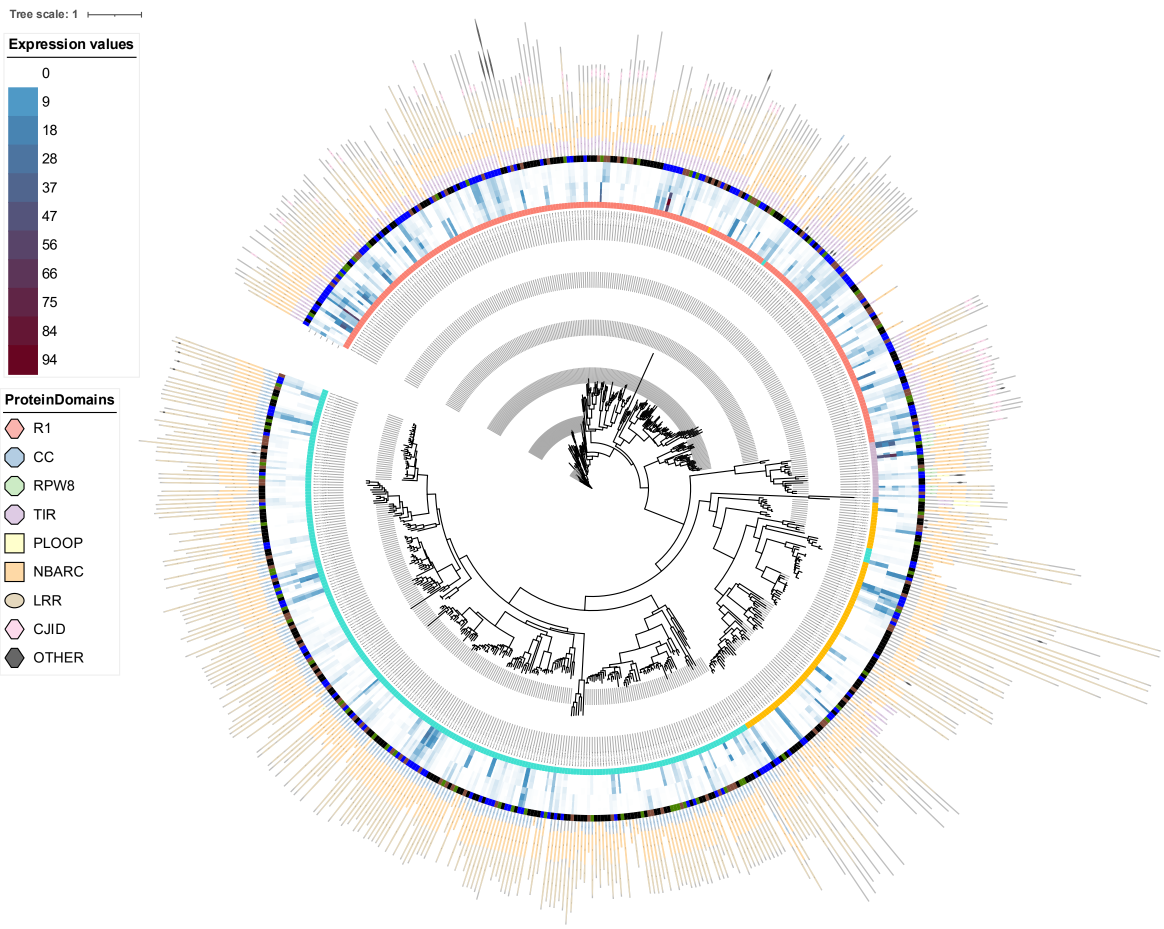

Tree scale: 1
Expression values
0
9
18
28
37
47
56
66
75
84
94
ProteinDomains
R1
CC
RPW8
TIR
PLOOP
NBARC
LRR
CJID
OTHER


F

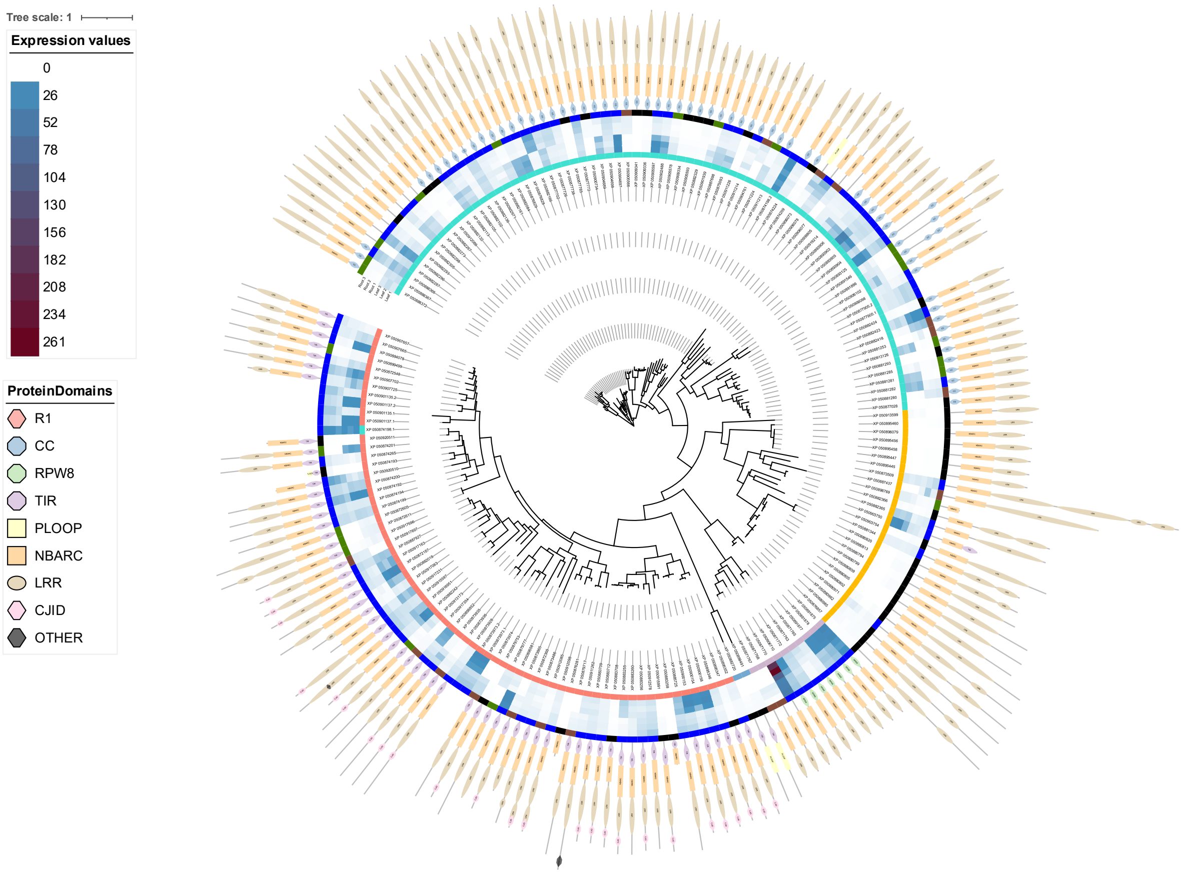

Tree scale: 1
Expression values
0
26
52
78
104
130
156
182
208
234
261
ProteinDomains
R1
CC
RPW8
TIR
PLOOP
NBARC
LRR
CJID
OTHER

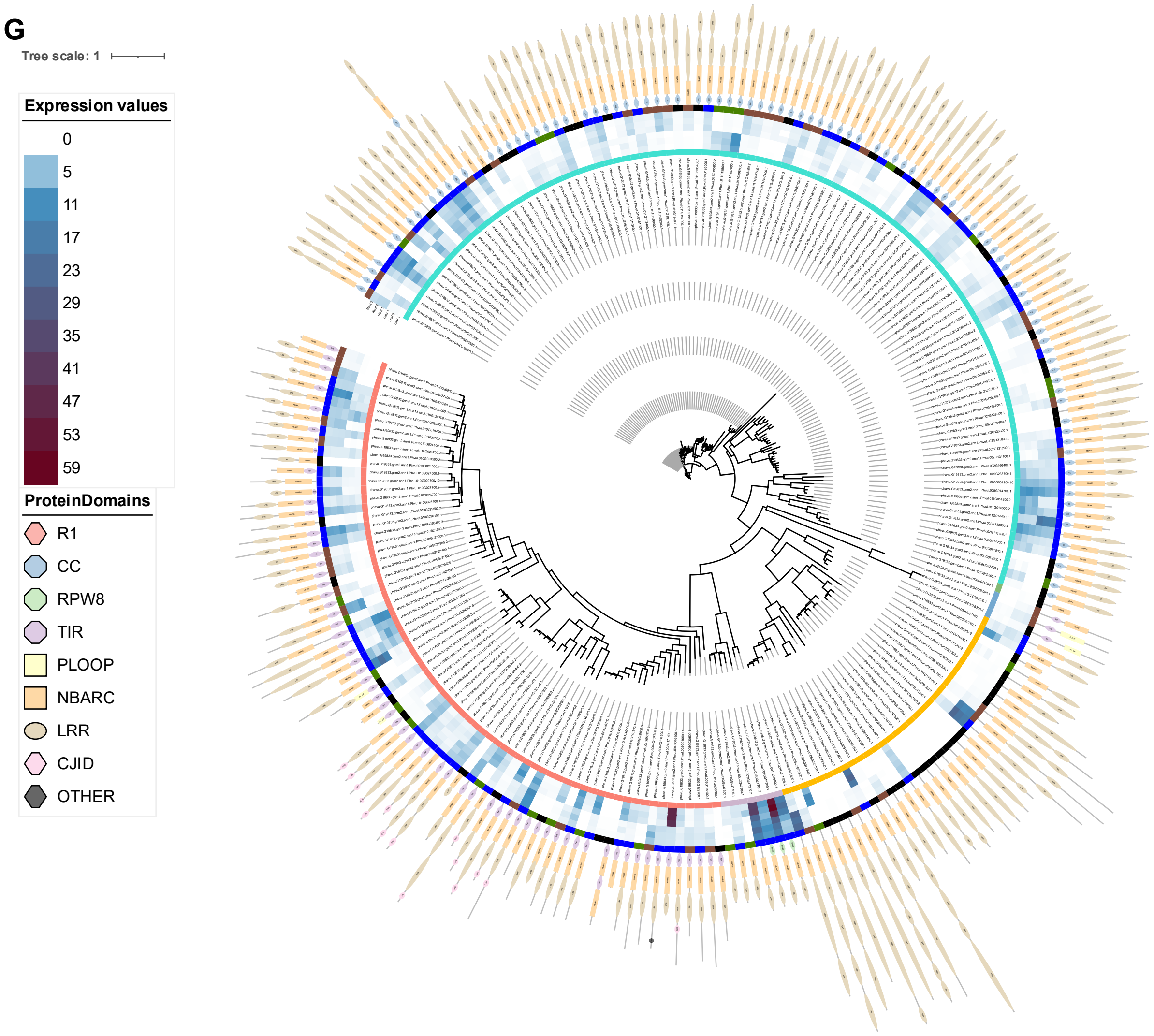


**Supplemental Figure 4. Species-specific analysis reveals patterns of NLR expression and tissue predominance.** Circular phylogenies displaying NLR expression for each of the seven species: **(A)** *Cicer arietinum*, **(B)** *Glycine max*, **(C)** *Lens culinaris*, **(D)** *Lathyrus sativus*, **(E)** *Medicago truncatula*, **(F)** *Pisum sativum*, and **(G)** *Phaseolus vulgaris*. For each plot, the concentric rings display, from innermost to outermost: **(i)** NLR class (CC-NLR: turquoise; TIR-NLR: salmon; $CC_{G10}$-NLR: yellow; $CC_{R}$-NLR: lilac; TNP: blue), **(ii)** TMM expression values across three leaf and three root biological replicates, and **(iii)** tissue-specificity classification (expressed in both tissues: blue; leaf-specific: green; root-specific: brown; not expressed: black).
